# ATP diffusional gradients are sufficient to maintain bioenergetic homeostasis in synaptic boutons lacking mitochondria

**DOI:** 10.1101/2022.12.09.519840

**Authors:** Ivan A. Kuznetsov, Andrey V. Kuznetsov

## Abstract

Previous work on mitochondrial distribution in axons has shown that approximately half of the presynaptic release sites do not contain mitochondria, raising the question of how the boutons that do not contain mitochondria are supplied with ATP. Here, we develop and apply a mathematical model to study this question. Specifically, we investigate whether diffusive transport of ATP is sufficient to support the exocytic functionality in synaptic boutons which lack mitochondria. Our results demonstrate that the difference in ATP concentration between a bouton containing a mitochondrion and a neighboring bouton lacking a mitochondrion is only approximately 0.4%, which is still 3.75 times larger than the ATP concentration minimally required to support synaptic vesicle release. This work therefore suggests that passive diffusion of ATP is sufficient to maintain the functionality of boutons which do not contain mitochondria.

## 1. Introduction

Mitochondria’s main purpose is to produce easily accessible chemical energy in the form of ATP [1,2]. In addition to stationary mitochondria, there are also anterogradely and retrogradely moving mitochondria [3–6]. ATP supply to energy demand sites, such as presynaptic boutons, was thought to be mainly provided by stationary mitochondria that dock in these boutons [7–9].

However, experimental research indicates that approximately half of the presynaptic release sites do not contain mitochondria [10–12]. An interesting question is how ATP is supplied into a bouton that lacks a mitochondrion. There are three competing hypotheses in the literature explaining how this can happen. 1) ATP is transported to boutons lacking mitochondria by diffusion [13]. 2) ATP is synthesized in boutons lacking a mitochondrion by glycolysis [13]. 3) ATP needs are partly met by ATP production in neighboring axons, dendrites, and glial cells [11,14]. The correct explanation can also be a combination of these hypotheses.

The main novelty of our paper is the explanation why axons do not need a stationary mitochondrion in every energy demand site. To explain this, we developed a mathematical model of diffusion-driven ATP transport from a bouton containing a mitochondrion to a bouton lacking a mitochondrion. Using the developed model, we investigated whether diffusion-driven transport of ATP is sufficient to maintain the ATP concentration in a bouton lacking a mitochondrion above the minimum ATP concentration required to support synaptic vesicle release. If diffusion is sufficiently fast, the ATP concentration will be approximately uniform; if it is slow, the ATP concentration will decay with the distance from a mitochondrion [11,14].

## 2. Materials and models

### 2.1. Equations governing ATP concentration in a single periodic unit control volume containing half of a mitochondrion

For the development of the model, we assumed that a mitochondrion is present in the center of every second varicosity (bouton) in the axon. This simplifying assumption of periodicity made it possible to define a periodic unit control volume containing half of a mitochondrion (the other half belongs to a neighboring periodic control volume), see Fig. 1. The width of axonal varicosity is 2*δ*; intervaricosity average spacing is *L*. Mitochondria are assumed to be located at every second varicosity. The distance between mitochondria is 2 *L* + 4*δ*.

**Fig. 1.**
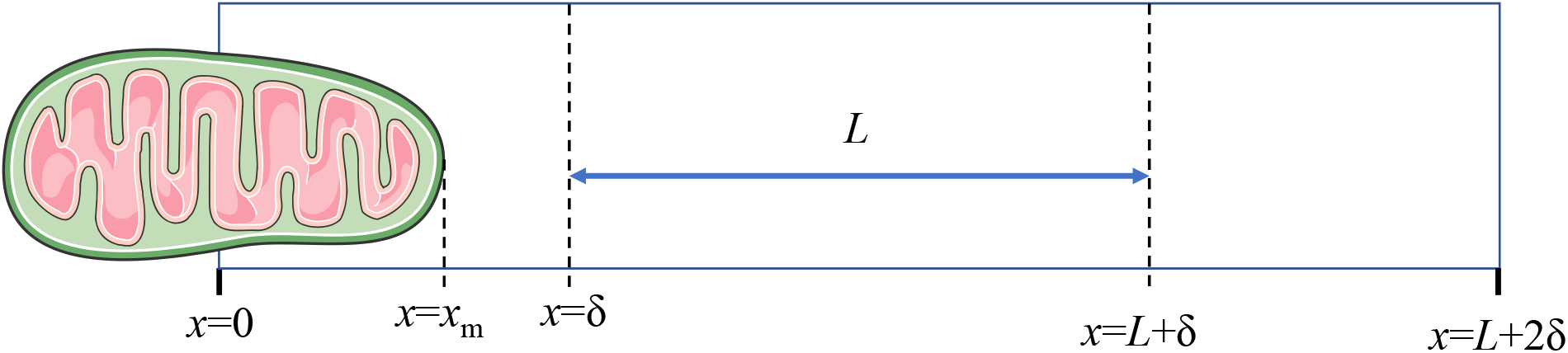
(a) Schematic diagram of a single periodic unit control volume containing half of a mitochondrion (the other half belongs to a neighboring periodic control volume). The width of axonal varicosity is 2*δ*; intervaricosity average spacing is *L*. Mitochondria are assumed to be located at every second varicosity. The distance between mitochondria is 2 *L* + 4*δ*. *Figure generated with the aid of servier medical art, licensed under a creative commons attribution 3.0 generic license. http://Smart.servier.com*.

Since mammalian cortex neurons spend a large amount of energy (~80%) on action potentials [15], ATP consumption is expected to oscillate over time. However, these oscillations are high-frequency, as cortical neurons fire ~0.16 times per second [15]. The timescale of ATP diffusion is much larger than the timescale of these spikes; therefore, in designing our model we use a time averaged value for the ATP consumption rate. Our model thus predicts time-averaged values of ATP concentrations.

In the portion of varicosity occupied by a mitochondrion (Fig. 1), the conservation of ATP molecules gives the following equation:

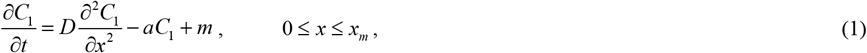

where *t* is the time and *x* is the linear coordinate along the axon.

In the portion of varicosity not occupied by a mitochondrion (Fig. 1), the ATP conservation is given by the following equation:

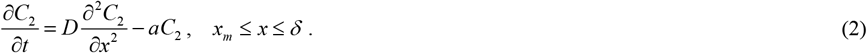

In the space between varicosities (also called intervaricosity spacing, Fig. 1), the ATP conservation results in the following equation:

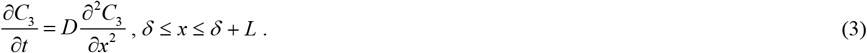

In an empty varicosity, not occupied by a mitochondrion (but still consuming ATP) (Fig. 1), the ATP conservation gives the following equation, which is similar to Eq. (2):

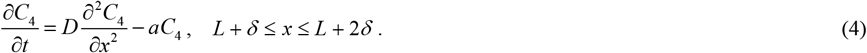

The only dependent variable utilized in the model is the linear concentration of ATP in the axon, *C*. Model parameters are summarized in Table 1.

**Table 1.**
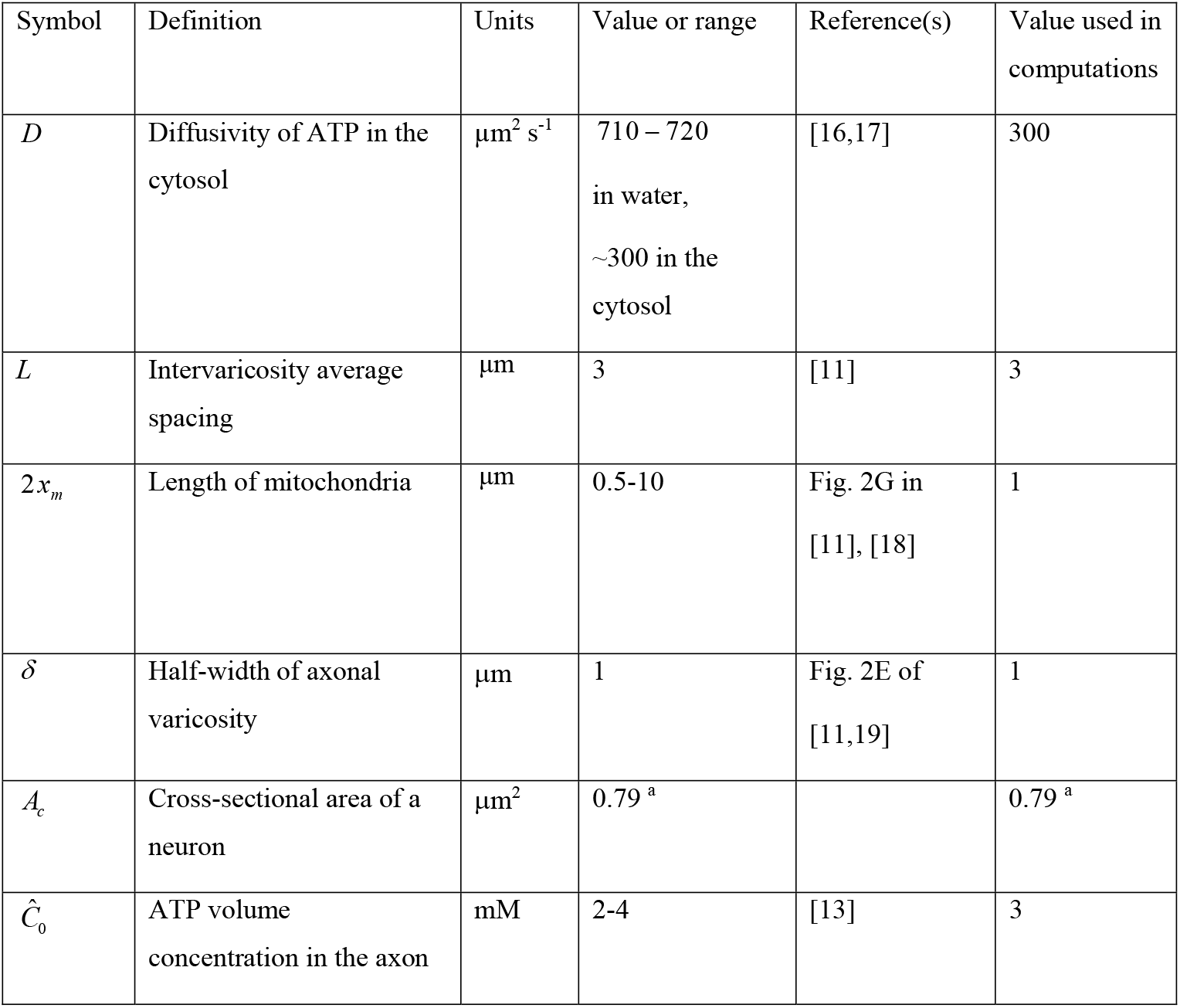
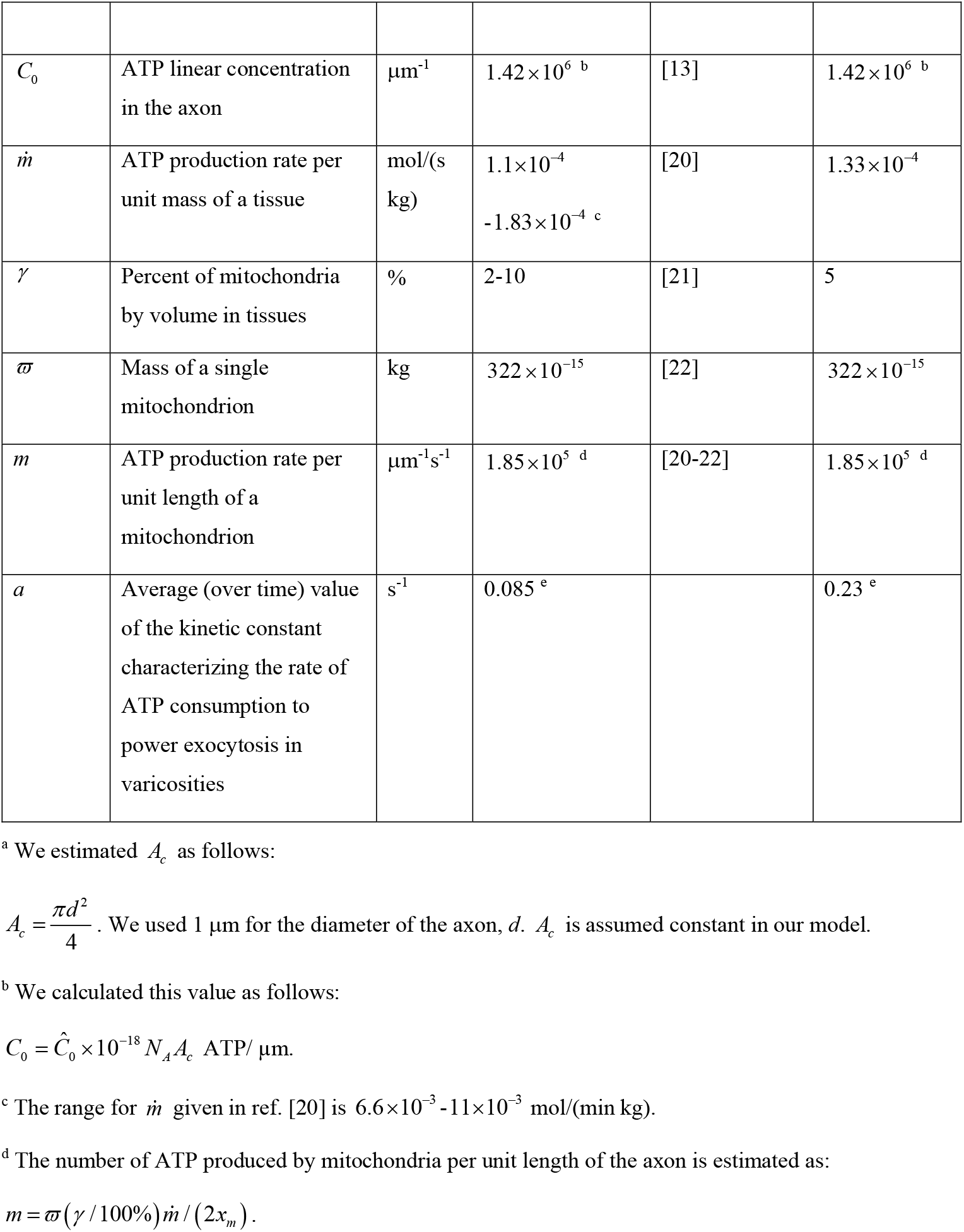
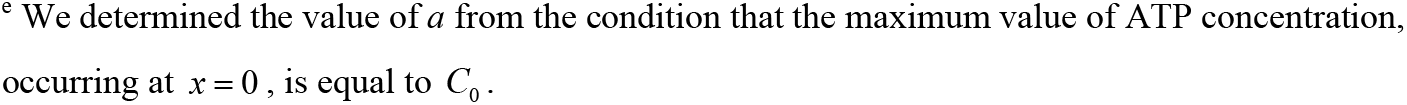
Parameters characterizing ATP transport and accumulation in the axon.

Eqs. (1)–(4) were solved at steady-state subject to the following boundary conditions:

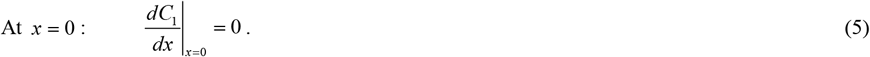

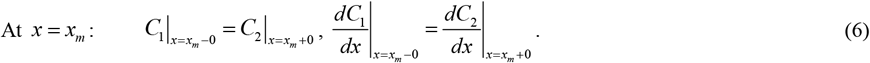

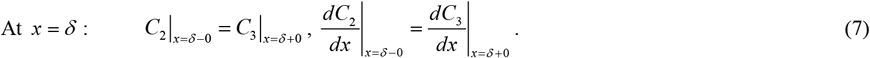

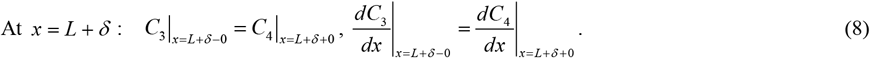

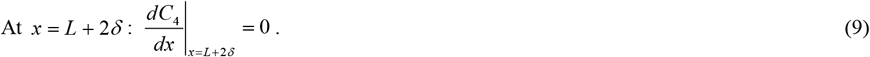

Note that the problem description is given in the transient formulation (see Eqs. (1)–(4)). This may be helpful for future extension of the present work.

## 3. Results

### 3.1. Distribution of ATP concentration between varicosities containing a mitochondrion and varicosities lacking a mitochondrion

The analytical solution of Eqs. (1)–(4) subject to boundary conditions (5)-(9) for the steady-state situation is given by Eqs. (S1)–(S4) in Supplementary Materials.

The maximum concentration of ATP occurs in the center of a bouton occupied by a mitochondrion (Fig. 2). Away from the mitochondrion, the ATP concentration decreases and reaches its minimum value in the center of varicosity that lacks a mitochondrion (Fig. 2). If the ATP concentration in the bouton containing a mitochondrion is 3 mM, the drop of the ATP concentration between the bouton containing a mitochondrion and a bouton lacking a mitochondrion is approximately 0.4%. An increase of intervaricosity spacing increases the ATP concentration drop between a varicosity containing a mitochondrion and a neighboring varicosity without a mitochondrion. According to [13], to sustain synaptic transmission, a minimum ATP concentration of approximately 0.8 mM needs to be maintained in boutons. This corresponds to

**Fig. 2.**
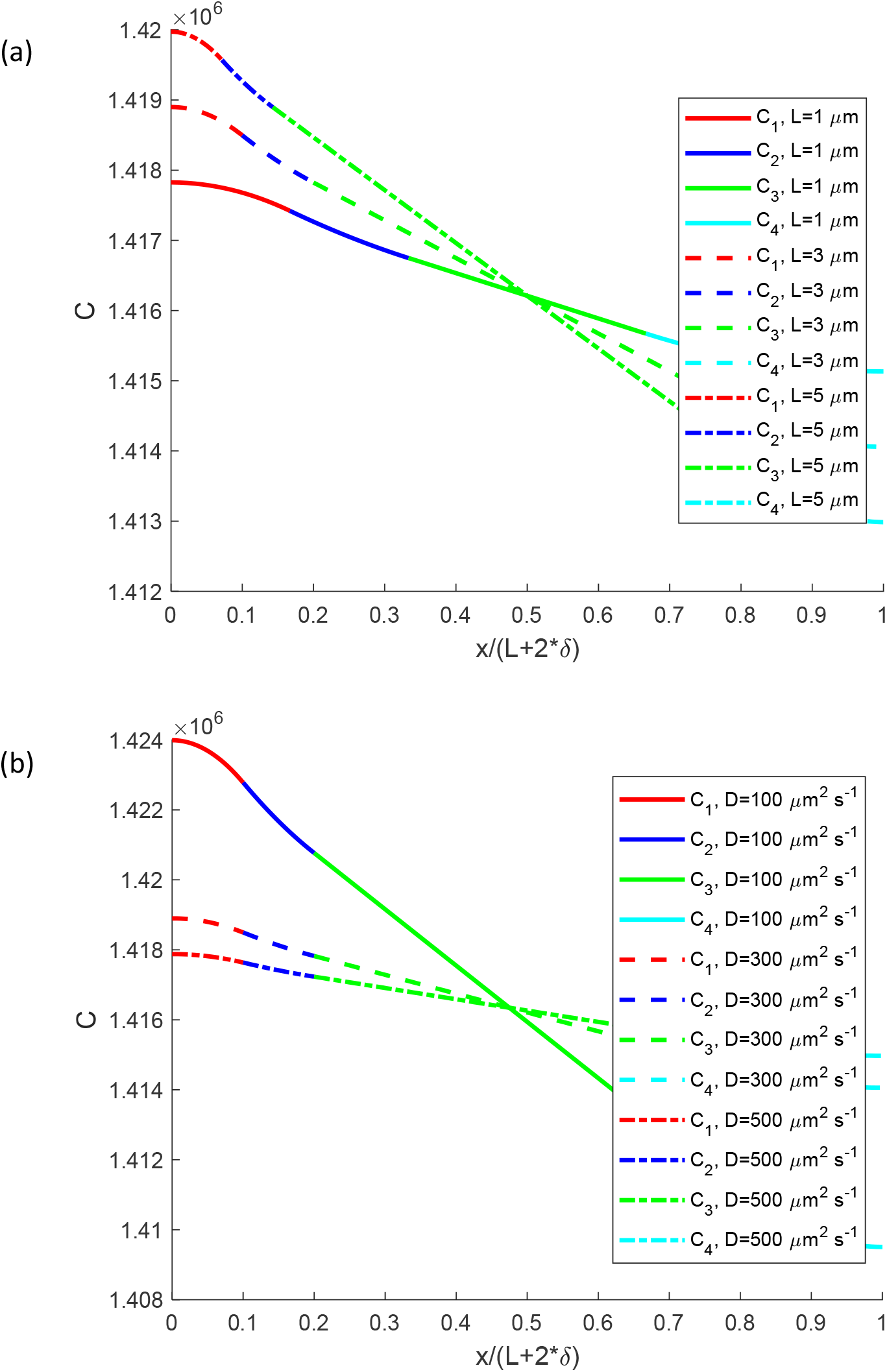
Linear concentration of ATP in the axon vs the distance from a mitochondrion for (a) different intervaricosity spacing; (b) different ATP diffusivities.

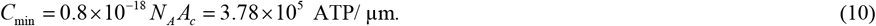

The concentration displayed in Fig. 2a is 3.75 times greater than the minimally required ATP concentration given by Eq. (10). This result is consistent with experimental observations reported in [13], which concluded that diffusive transport of ATP is sufficiently fast for ATP distribution to be uniform, even in boutons lacking mitochondria.

Computations with a smaller value of ATP diffusivity predict a larger drop between the ATP concentration in a bouton containing a mitochondrion and a bouton lacking a mitochondrion (approximately 1% for *D* = 100 µm^2^ s^−1^, Fig. 2b). Computations with a greater value of ATP diffusivity predict a more uniform ATP distribution (the concentration drop is approximately 0.14% for *D* = 500 µm^2^ s^−1^, Fig. 2b).

### 3.2. Sensitivity of the concentration of ATP to ATP diffusivity and intervaricosity spacing

This investigation was performed by computing local sensitivity coefficients, which are first-order partial derivatives of the observables with respect to model parameters [23–26]. For example, the sensitivity coefficient of the ATP concentration to ATP diffusivity *D* at steady-state (ss) was calculated as follows:

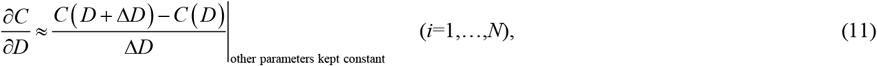

where Δ*D* =10^−2^ *D* (we tested the accuracy by using various step sizes).

The sensitivity coefficient was non-dimensionalized to make it independent of the magnitude of the parameter whose sensitivity was tested [24,27], as follows (for example):

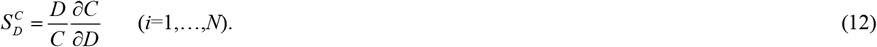

The dimensionless sensitivity of ATP concentration to the ATP diffusivity is positive and increases with the distance from the mitochondrion (Fig. 3a). This is because the increase of ATP diffusivity increases the rate of diffusion-driven transport of ATP, making the ATP concentration in the axon more uniform, closer to that in the region occupied by the mitochondrion.

**Fig. 3.**
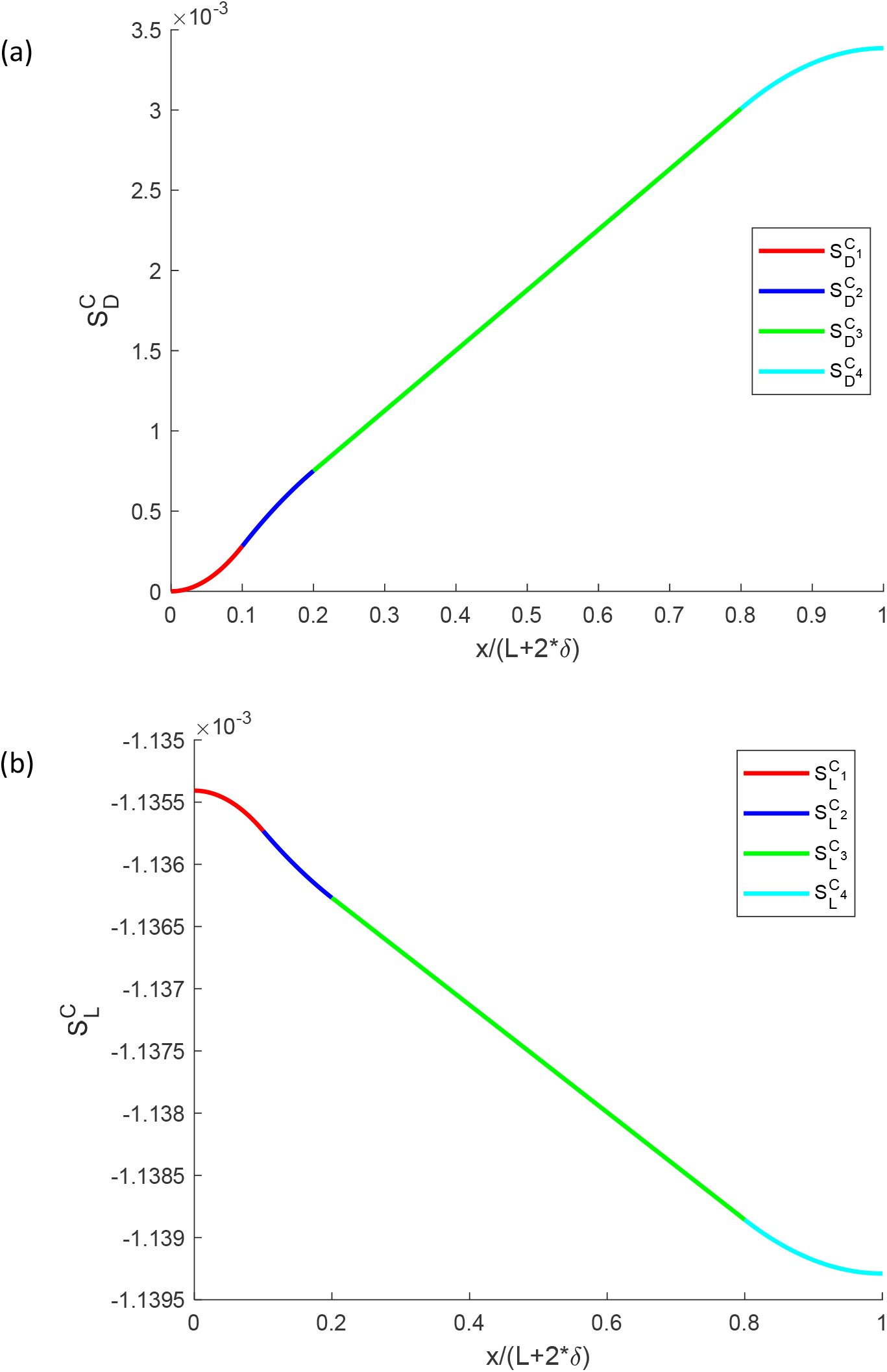
(a) Dimensionless sensitivity of the ATP concentration to the ATP diffusivity. Computations were performed with Δ*D* = 10^−2^ *D*. A close result was obtained for Δ*D* = 10^−1^ *D*. (b) Dimensionless sensitivity of the ATP concentration to the intervaricosity spacing. Computations were performed with Δ*L* = 10^−2^ *L*. A close result was obtained for Δ*L* = 10^−1^ *L*. Sensitivities are analyzed around *D* = 300 µm^2^ s^−1^, *L* = 3 μm.

The dimensionless sensitivity of ATP concentration to the intervaricosity spacing is negative and becomes more negative with an increase in the intervaricosity spacing (Fig. 3b). This is because diffusion becomes a less efficient transport mechanism with an increase in the distance that ATP must be transported.

## 4. Discussion, limitations of the model, and future directions

We investigated the variation in ATP concentration between a bouton containing a mitochondrion and a bouton lacking a mitochondrion. An analytical solution of equations governing the ATP diffusion is obtained. We established that in the case of a mitochondrion being present only in every second bouton, the drop in the ATP concentration between a bouton containing a mitochondrion and a bouton lacking a mitochondrion is only 0.4%. Therefore the ATP concentration in the axon is practically uniform due to the large diffusivity of ATP, thus explaining why axons do not need a stationary mitochondrion in every energy demand site. Thus the results of our paper support the conclusions of the experimental work [13] that the ATP diffusion helps maintain a sufficient level of ATP derived from the mitochondria to ensure proper synaptic transmission, even in synaptic boutons that do not contain mitochondria.

The sensitivity of the ATP concentration to the ATP diffusivity is positive, which means that an increase in ATP diffusivity makes the ATP concentration in the axon more uniform. Conversely, the sensitivity of ATP concentration to the intervaricosity spacing is negative, which means that an increase in intervaricosity spacing makes the ATP concentration less uniform.

One of the limitations of the proposed model is the assumption that the ATP consumption rate is directly proportional to ATP concentration (Eqs. (1), (2), and (4)). Better ATP consumption models need to be incorporated into future research. Future work should also investigate the 1-D transient problem given Eqs. (1)–(9) (rather than steady-state solved in this paper).

## Acknowledgment

IAK acknowledges the fellowship support of the Paul and Daisy Soros Fellowship for New Americans and the NIH/National Institute of Mental Health (NIMH) Ruth L. Kirchstein NRSA (F30 MH122076-01). AVK acknowledges the support of the National Science Foundation (award CBET-2042834) and the Alexander von Humboldt Foundation through the Humboldt Research Award.

## Ethical Statement

None.

## Supplementary Materials

### S1. Analytical solutions of governing equations

Analytical solutions of Eqs. (1)–(4) with boundary conditions (5)-(9) at steady-state are given by the following equations:

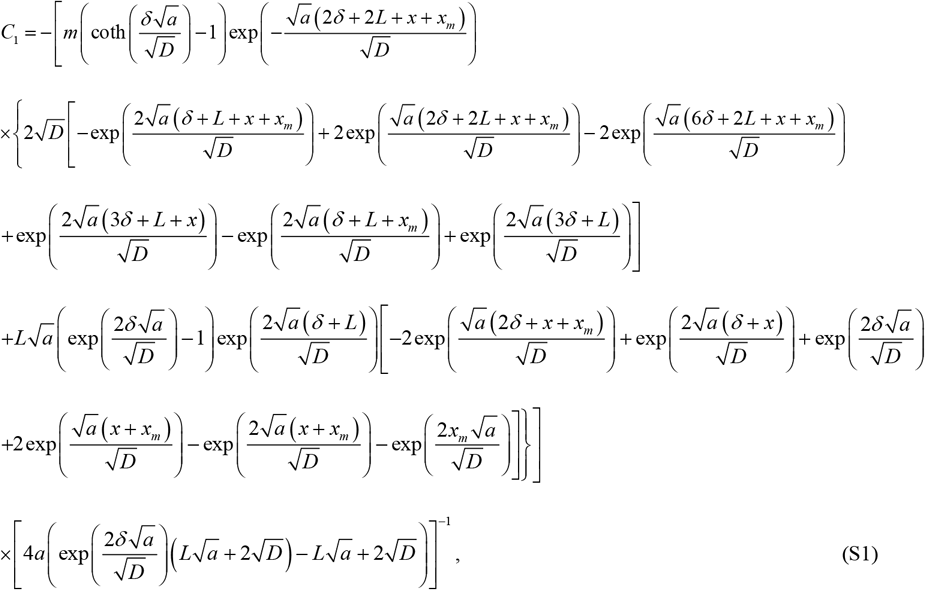

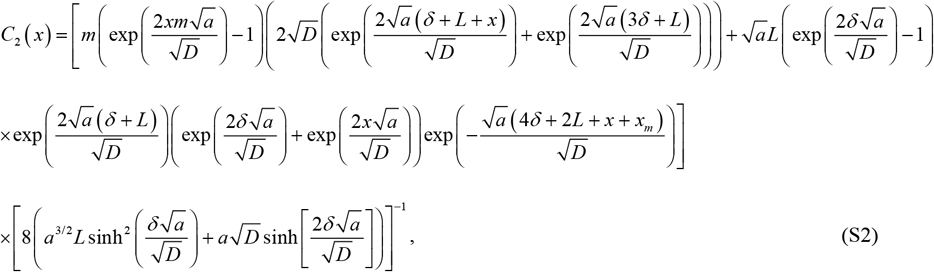

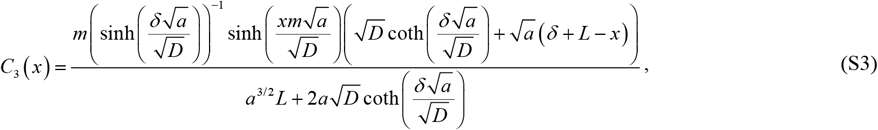

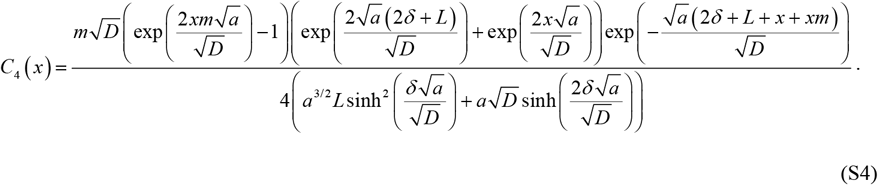

